# Targeted spectroscopy in the eye fundus

**DOI:** 10.1101/2023.04.27.538643

**Authors:** Nicolas Lapointe, Cléophace Akitegetse, Jasmine Poirier, Maxime Picard, Patrick Sauvageau, Dominic Sauvageau

## Abstract

**Significance:** The assessment of biomarkers in the eye is rapidly gaining traction for the screening, diagnosis and monitoring of ocular and neurological diseases. Targeted ocular spectroscopy is a new technology that enables the user to concurrently image the eye fundus and acquire high quality spectra from a targeted region –1.5 degrees– within the imaged area. The combination of imaging and high-sensitivity spectroscopy provides structural, compositional, and functional information of selected regions of the eye fundus. This opens the door to new, non-invasive approaches to the detection of biomarkers in the eye.

**Aim:** The aim of this study was to demonstrate the multi-modal functionality and validation of the targeted ocular spectroscopy developed. This was done in vitro, using a reference target and a model eye, and in vivo.

**Approach:** Images and spectra from different regions of a reference target and a model eye were acquired and analyzed to validate the system. The same eye model was used to obtain fluorescence images and spectra, highlighting the capability of the system to also perform targeted ocular fluorescence spectroscopy. Subsequently, in vivo imaging and diffuse reflectance spectra were acquired to assess blood oxygen saturation in the optic nerve head and the parafovea of healthy subjects.

*Results:* Tests conducted with the reference target showed that spectral analysis could be accurately performed within specific areas of the imaging space. Moving to the model eye, distinct spectral signatures were observed for the targeted spectral analysis done in the optic disc, the retina and the macula, consistent with the variations in tissue composition and functions between these regions mimicked by the model eye. Further, it was shown that the targeted spectral analysis could also be performed in a fluorescence mode to distinguish various fluorophores present within the imaging space. Finally, in vivo ocular oximetry experiments performed in the optic nerve head and parafovea of healthy patients showed significant differences in blood oxygen saturation between these regions (p = 0.004).

*Conclusions:* Enabling non-invasive, sensitive diffuse reflectance and fluorescence spectroscopy in specific regions of the eye fundus opens the door to a whole new range of monitoring and diagnostic capabilities, from assessment of oxygenation in glaucoma and diabetic retinopathy to photo-oxidation and photo-degradation in age-related macular degeneration.

## 1 Introduction

Ocular diseases generally involve specific structural and functional changes in the eye fundus. For example, a progressive degeneration of ganglion cells characterizes glaucoma [1], accumulation of drusen and auto-fluorescent pigments are observed in age-related macular degeneration (AMD) [2], and micro-aneurysms, retinal hemorrhages and vascular abnormalities are associated with diabetic retinopathy (DR) [3]. Morphological and functional changes in the eye fundus are, however, not strictly restricted to vision-related diseases. Recent studies have shown that certain neurological diseases, such as Parkinson and Alzheimer, lead to observable changes in the retina, such as thinning of the retinal nerve fiber layer (RNFL) and changes in hemodynamics [4]–[10].

Spectral analysis of light reflected or emitted by the retinal tissue can provide valuable information on changes in the retina that cannot be uncovered through typical fundus color imaging or optical coherence tomography (OCT). This approach has thus great potential as a complement to current assessment tools for a number of research and clinical applications. For instance, accumulation of lipofuscin in the retinal pigment epithelium (RPE) causes fundus autofluorescence [11], [12], structural changes in RNFL impact retinal reflectance [13], [14], while the absorption spectrum of blood varies according to the ratio of its content in oxygenated and reduced hemoglobin [15]. Such localized changes have different outcomes that impact the optical properties of retinal tissues, which can be measured through diffuse reflectance spectroscopy.

Diffuse reflectance spectroscopy has been the subject of multiple studies over the last decades. For example, in 1986, Van Norren and Tiemeijer used a retinal densitometer to measure the spectral reflectance of the optic disc, the peripheral retina and the fovea [16] with an angular resolution of 2.5 degrees. The authors were able to establish that reflectance was higher in the red range than in the blue range and came up with a model to explain their measurements. Delori and Pflibsen used a modified fundus camera to acquire retinal reflectance spectra from 450 nm to 800 nm. They achieved an angular resolution varying from 1 to 4 degrees by placing an aperture in the illumination path to limit the illuminated area [17]. A grating monochromator and a camera were used for spectral acquisition, and a fixation target was used to change the measurement site. Later, Delori developed a system allowing the acquisition of both the intrinsic fluorescence spectrum and the reflectance of the eye fundus in the spectral range of 500 nm to 800 nm using a modified ocular fluorometer [17]. To achieve this, an aperture in the image plane of the retina was used to define the sampling area. Hammer *et al.* used a slit-shaped stop in an image plane of the eye fundus to acquire reflectance spectra of horizontal areas [18]. Diaconu and Vucea *et al.* proposed a system using a holed-mirror located in an image plane to reflect light coming from the fundus to a camera with the same intent [19], [20]. The light that passed through the small hole was acquired by a spectrometer, and thus the spectral sampling area was the center of the image, where a black region was visible due to the hole in the image plane.

Although allowing the acquisition of a localized spectral signal in the eye fundus, the aforementioned techniques lack flexibility in the sense that the acquisition area is only adjusted by the fixation of the subject, which can be laborious when fine positioning is required.

Moreover, except for the method developed by Diaconu [20], spectral acquisition could not be done while simultaneously observing the acquisition area or imaging the eye fundus. Under these conditions, the actual position of the acquisition area is difficult to determine.

Hyperspectral imaging systems allow for the acquisition of eye fundus images with several color channels, such that each pixel contains spectral information. Some of these systems consist of retinal cameras where the transmitted wavelength band is controlled by a tunable filter applied either in imaging or illumination to construct the hyperspectral cube [21], [22]. Other technologies use a diffractive grating to separate colors, and each color is collected by a dedicated area of the detector [23], [24]. However, all these systems face a compromise between spectral resolution, acquisition speed, and spatial resolution. Image registration is also required most of the time. Hence, at least at this point in time, these systems are not as sensitive as technologies based on spectrometers. Desjardins *et al.* reported hyperspectral imaging in under 3 sec for a FOV of 30 degrees and a wavelength range of 500 to 600 nm with 2 nm and 5 nm steps [22]. In a more recent study, the same team achieved an overall acquisition time of under 1 sec for a range of 450 to 900 nm in steps of 5 nm [25]. These factors contribute to a limitation in the sensitivity to some biomarkers and the response time of these systems.

Visible-light optical coherence tomography (VIS-OCT) is another technology that enables both anatomical and functional imaging. Typical OCT devices use near-infrared (NIR) light, but using a super-continuum light source in the visible spectral range enables higher resolution and acquisitions with absorption contrast information. Different groups have demonstrated the ability of VIS-OCT to extract metabolic information from visible light spectra. Pi *et al*. measured, quantitatively, retinal blood oxygen saturation *in vitro* and in rats [26]. Recently, Song *et al*. have shown that the reflectance signal from VIS-OCT could better distinguish glaucoma subjects from normal eyes than thickness measurements from OCT [27]. However, the acquisition time was 2.6 sec at 50 kHz A-line speed and the laser power at the pupil was 0.25 mW for a wavelength range of 545 to 580 nm. In another paper, Song *et al.* reported VIS-OCT acquisitions of less than 6 sec for a field of view (FOV) of 3 mm × 7.8 mm and a wavelength range of 545 to 580 nm [28]. This requires the patient to have a stable fixation while being subjected to high irradiance for the whole duration of the acquisition, which is a major drawback and represents a challenge for the clinical adoption of this technology.

Considering these technologies, their limitations and the structural, functional and compositional heterogeneity of the eye fundus, there remains an unmet need for the spectral assessment of specific regions of the eye fundus. We describe, here, a system that allows the visualization of the eye fundus while simultaneously acquiring full visible diffuse reflectance or fluorescence spectra from a selected location of the eye fundus. The user can target any visible location within the eye fundus region being imaged without any realignment or change of fixation target, while continuously receiving spectral information of the targeted sampled area. Such assessments can potentially provide important information for the detection, diagnosis and/or monitoring of ocular and other diseases.

## 2 Materials and Methods

### 2.1 Targeted optical spectroscopy

The optical layout used for concurrent imaging and spectroscopy is based on the Zilia Ocular platform (Figure 1). Its core is similar to that of a typical fundus camera. Light from two 5000K white light emitting diodes (LEDs) (YJ-BC-3030-G04, Yujileds, Beijing, China) and two NIR LEDs centered at 785 nm (SST-10-FR-B90-H730, Luminus, Sunnyvale, USA) is magnified through a custom-made 4-f system and reflected by a holed mirror through the pupil using an objective lens, as can be seen in the illumination path of Figure 1. The white light is filtered via a long-pass 495-nm filter (AT495lp, Chroma). The user can choose to use the white or NIR LEDs for fundus illumination. In order to reduce reflections from the cornea, a mask is placed in a plane conjugate to the pupil, while reflections on the objective lens are minimized by the use of a black dot in a plane conjugate to the back surface of the objective lens. The light reflected on the eye fundus is collected by the objective lens, passes through the hole of the holed mirror and is projected to a CMOS camera via another custom-made 4-f system. A sputter-coated, non polarizing beam splitter (CHROMA, PN 21020, 80/20) is placed at the Fourier plane of the imaging system to project a fraction of the light oriented toward the camera in the direction of a multimode circulator (Castor Optics Inc., WMC2L1-C). This fraction of light is then sent through a group of lenses and is collected and analyzed by a spectrometer (C13555MA, Hamamatsu corporation, Hamamatsu City, Japan) connected to the circulator.

**Figure 1.**
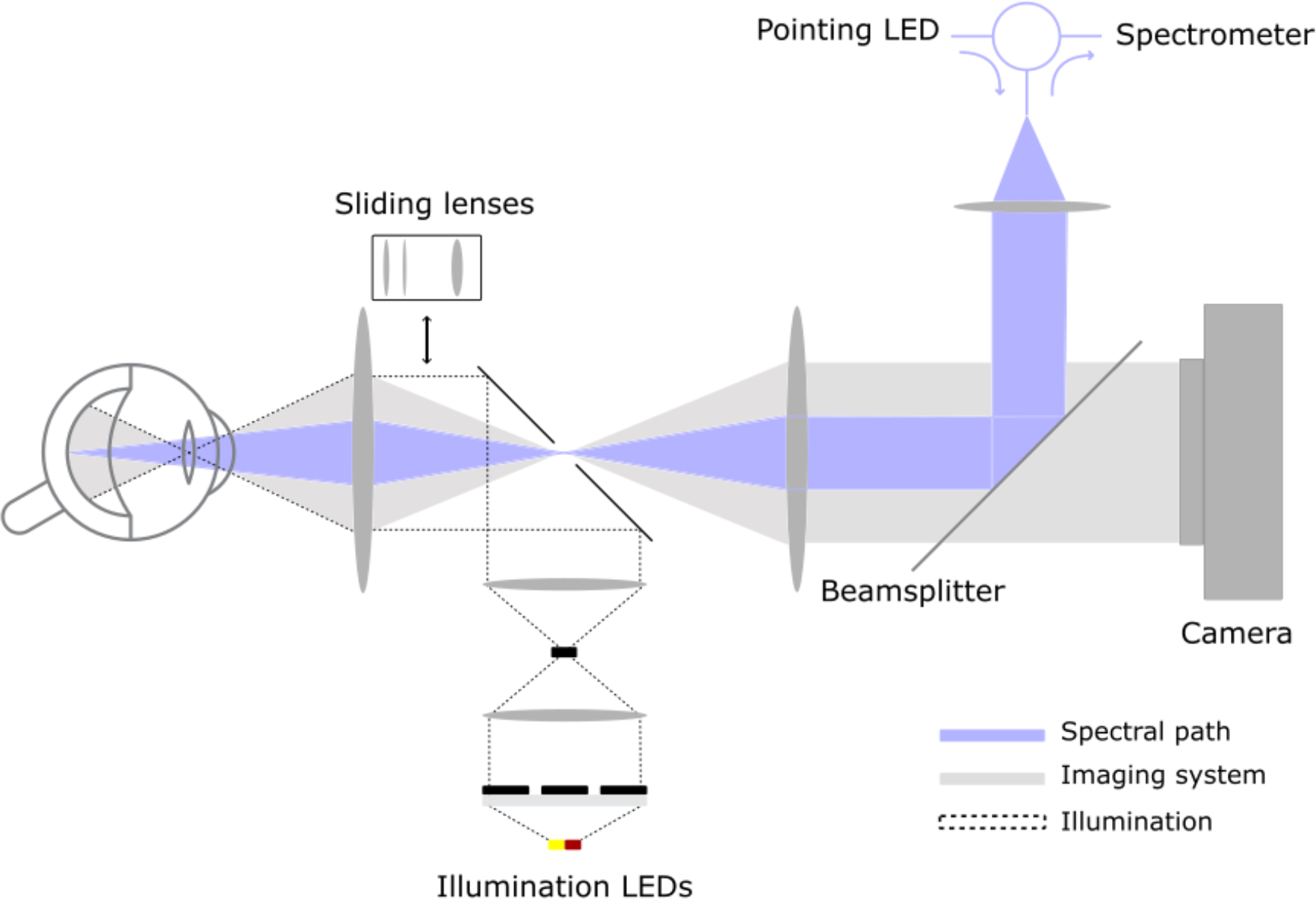
Targeted spectroscopy optical layout. Illumination LEDs illuminate the retina uniformly. Reflected light is directed partially toward the camera and the spectrometer. The Pointing LED is projected onto the eye fundus to identify the region of spectral acquisition (ROSA). Light going into the spectrometer comes from the ROSA, as it shares the same optical path as the pointing LED.

In order to identify the region of spectral acquisition (ROSA) – an area of 1.5 degree on the eye fundus for which spectral analysis is performed –, a 730-nm LED, referred to as the pointing LED, (SST-10-FR-B90-H730, Luminus, Sunnyvale, USA) is connected to the remaining port of the multimode circulator. This configuration allows the light collected by the spectrometer and that emitted by the pointing LED to share the same optical path, from the exit of the circulator to the eye fundus. Thus, when the pointing LED is activated, it illuminates the exact position of the ROSA and the camera can capture an image of its location. Two infrared LEDs (VSMY98145DS, Vishay, USA) placed on either side of the objective lens illuminate the outside of the eye, while a group of lenses installed on a linear actuator (sliding lenses in Figure 1) can be inserted behind the objective lens thus enabling corneal imaging.

A two-step acquisition sequence has been developed to combine imaging and targeted spectroscopy. First off, the illumination LED is turned on, illuminating the eye fundus and enabling simultaneous acquisition of an image by the camera and of a diffuse reflectance spectrum (DRS) by the spectrometer (Figure 2a and 2b). Then, the illumination LED is turned off and an image is acquired with only the pointing LED turned on (Figure 2c). This enables the identification and segmentation of the ROSA, while preventing cross-talk in the optical fiber.

**Figure 2.**
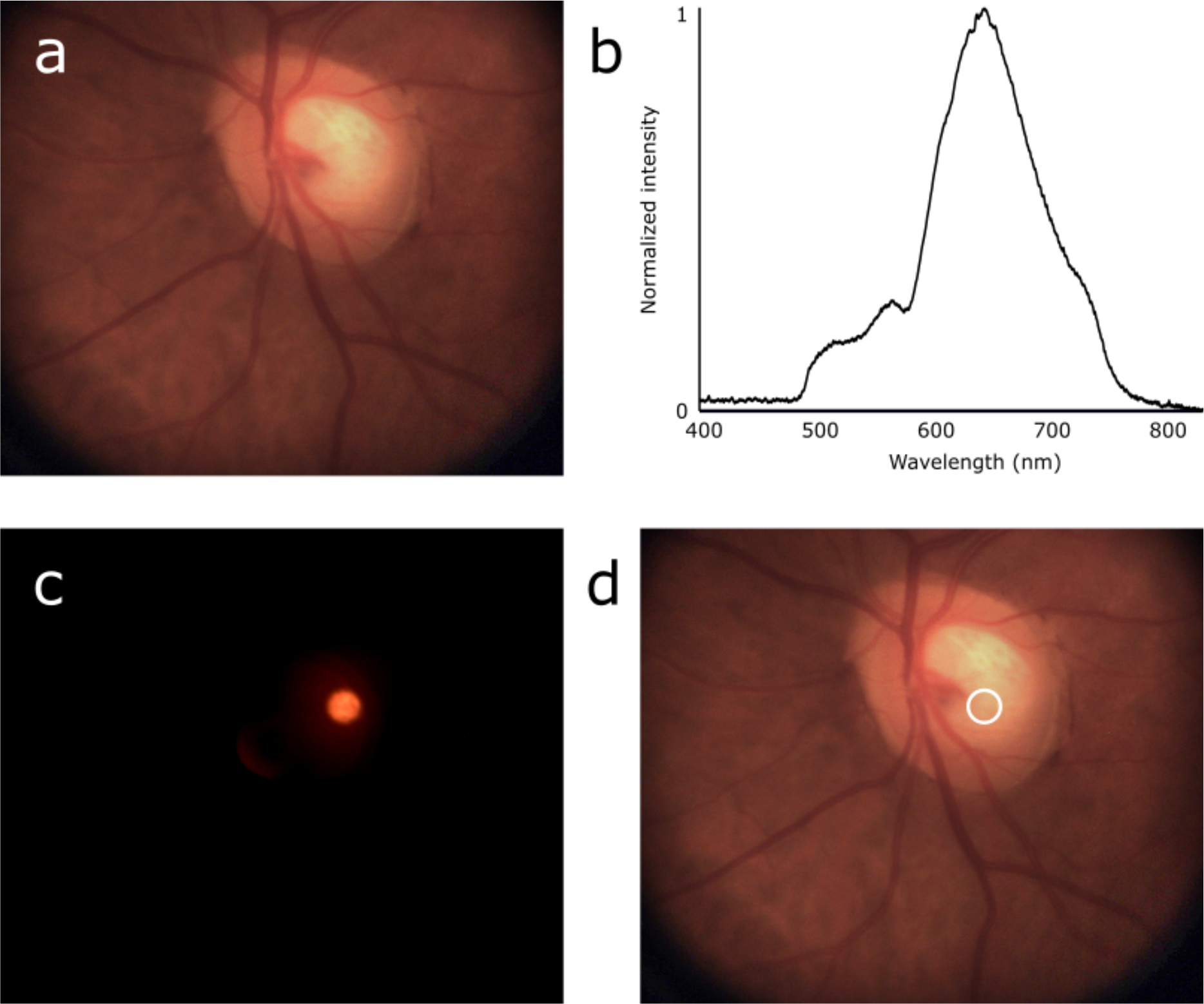
Principles of operation of targeted spectroscopy, enabling continuous, real-time imaging and spectral acquisitions. a) A white LED is used to illuminate the eye fundus, an image is acquired by the camera. b) A spectrum is simultaneously acquired by the spectrometer. c) The white LED is turned off and the camera acquires an image of the pointing LED projected onto the eye fundus, thus defining the ROSA. d) An image of the eye fundus overlaid with the location of the ROSA is shown to the user.

The segmentation of the ROSA image is used to identify the location of the DRS acquisition area (Figure 2d).

### 2.2 Reference Target

To demonstrate the ability of the system to acquire a spectrum from a selected region, an ultra-high-definition screen (HP V28 4K, 3840 pixels by 2160 pixels and pixel pitch of 0.116 mm) was used to display a reference target. The target displayed on the screen consisted of an 8 x 8 grid with 8 different colors (Figure 4a). Each chosen color consisted of an RGB color code, with each component at either 0 or maximum intensity. The fundus camera with a 150-mm lens (Edmund optics, PN #47-352) at its entrance pupil was positioned in front of the screen (Figure S1)

### 2.3 Model Eye

To validate the ability of the system to perform DRS acquisitions from the eye fundus, an imaging Eye Model (OEMI-7, Optical Instruments, Bellevue, Washington) with a 7-mm pupil was used (Figure 3). The OEMI-7 eye model is designed to accurately simulate the human eye; it includes an anterior chamber and crystalline lens. It displays a representation of the macula, a foreign body, the optic disc and blood vessels (Figure 3a). Moreover, the model has fluorescent patterns, allowing for simulated fluorescence imaging (Figure 3b).

**Figure 3.**
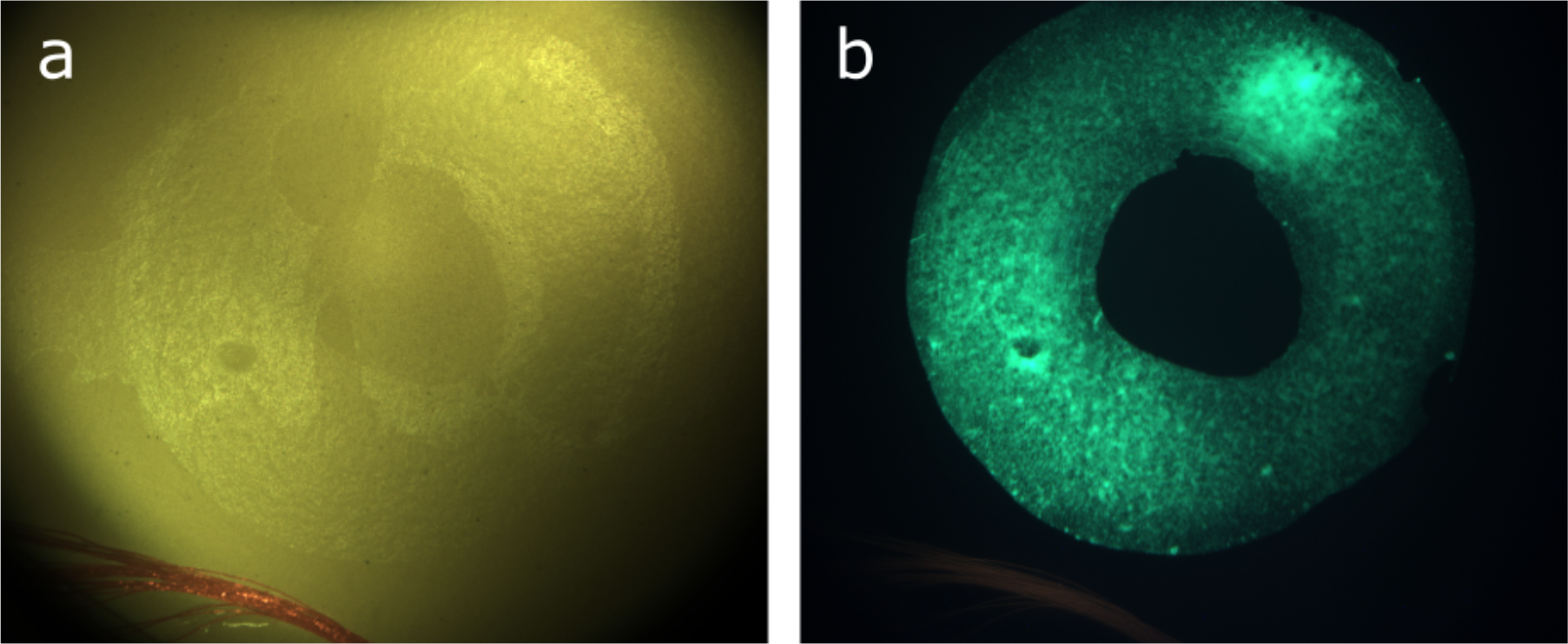
Ocular Imaging Eye Model OEMI-7. a) White reflectance image. b) Fluorescence imaging.

### 2.4 Fluorescence Acquisition

Slight modifications of the optical design enabled the assessment of fluorescence through imaging and spectroscopy. A band-pass filter (AT460/50, Chroma Technology, Bellows Falls, USA) was inserted in the illumination path to select the excitation illumination for green fluorescence. A long-pass filter (AT495LP, Chroma Technology, Bellows Falls, USA) was also added to the system, behind the holed mirror. This filter enabled imaging of the fluorescent emitted light. The 730-nm LED used for pointing was replaced by a 450-nm laser diode. The spectrometer integration time was set at 15 ms. The power for the excitation source was set to 1 mW.

### 2.5 Spectral analysis

Each acquired raw spectrum went through the same processing steps. This consisted of removing the signal coming from the environment, then removing the spectrometer background, and normalizing the spectrum by the standard deviation of the raw signal from 420 nm to 700 nm (effective range of the white color spectrum) to correct for differences in signal intensity. The background was computed by averaging the spectral intensity between 350 nm and 400 nm, where the only spectral contribution was electrical noise from the spectrometer, since no UV light was present in the signal.

### 2.6 Human subjects

Eight non-smoking, healthy adults (mean age: 32 years old, range: 27–35 years, 4 males/4 females) were enrolled. They presented no systemic disease and took no medication. For all subjects, ophthalmic examination – including slit lamp biomicroscopy, measurement of visual acuity, objective refraction, Goldmann tonometry and funduscopic examination – produced normal findings. Exclusion criteria were ocular disease, acute infection, known diabetes mellitus, epilepsy, history of systemic hypertension or abnormal clinical optic disc appearance. After the examination, the right eye was dilated using tropicamide 1% and phenylephrine 2.5%. A 20-min acclimation period followed. The protocol of the study followed the guidelines of the Declaration of Helsinki. Signed informed consent was obtained from all subjects after the nature of the study was explained, and all participants were informed of their right to withdraw from experimentation at any point in the study.

### 2.7 Data processing

The spectra acquired were corrected using the method described in section 2.5. Data acquired from human subjects underwent additional processing to derive oxygen saturation (StO2). The process for correcting spectra and calculating oxygen saturation has been described elsewhere [29].

## 3 Results

### 3.1 Targeted spectroscopy

Demonstration of the ability of the system to acquire a spectrum from a specific targeted region within the imaging field of view was performed using the reference target described above. Data were acquired from 5 different locations in the reference target and spectra were compared.

Regions were selected by displacing the red LED projection (corresponding to the ROSA) on the screen. The white circles in Figure 4 represent the different regions where the ROSA was positioned.

**Figure 4.**
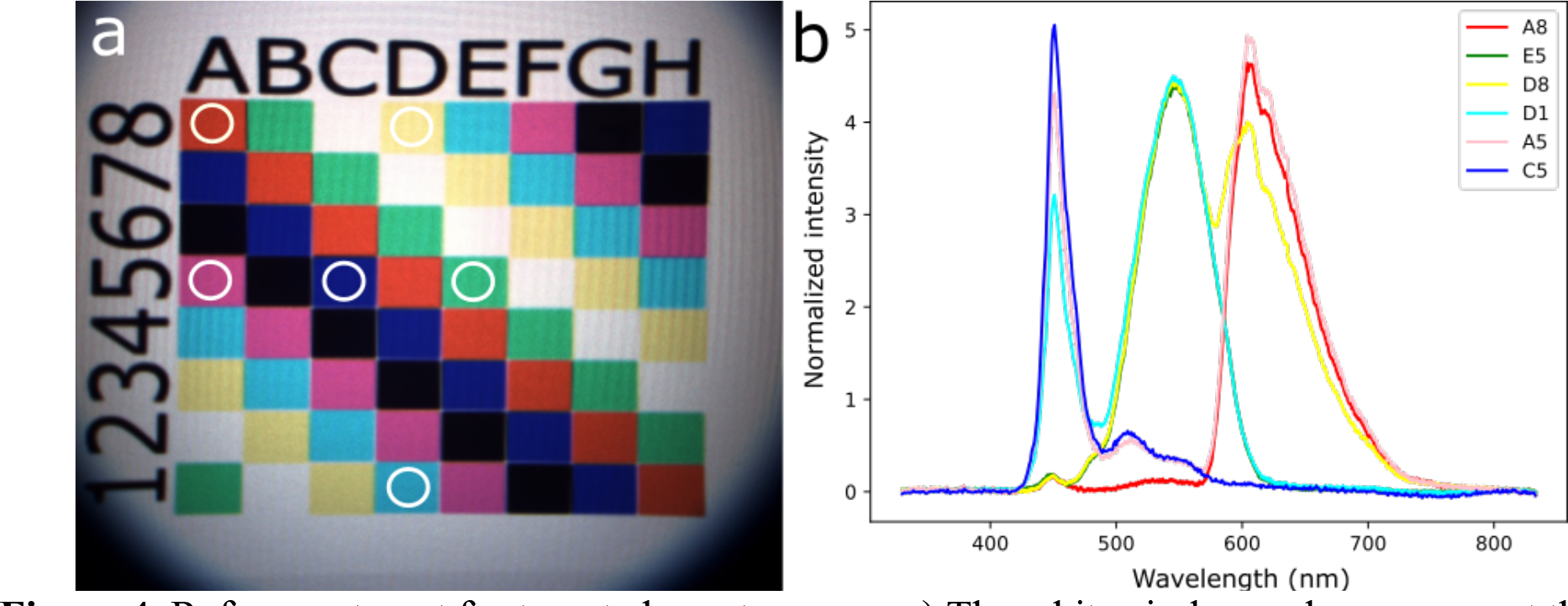
Reference target for targeted spectroscopy. a) The white circle overlays represent the different projections of the pointing LED for the regions tested. b) The spectra associated with each region tested.

To confirm that the region illuminated by the pointing LED precisely matched the ROSA, spectra were acquired on two concentric circles of different colors (red and blue) and dimensions (Figure 5). The inner circle diameter was set to different values to represent limit cases of the ROSA diameter. The ROSA was centered on the internal circle. The contribution of the red peak, at 625 nm, was only detected in the third and fourth acquisitions, where the ROSA slightly overlapped with the red color, indicating that the measure was localized within the area defined by the pointing light.

**Figure 5.**
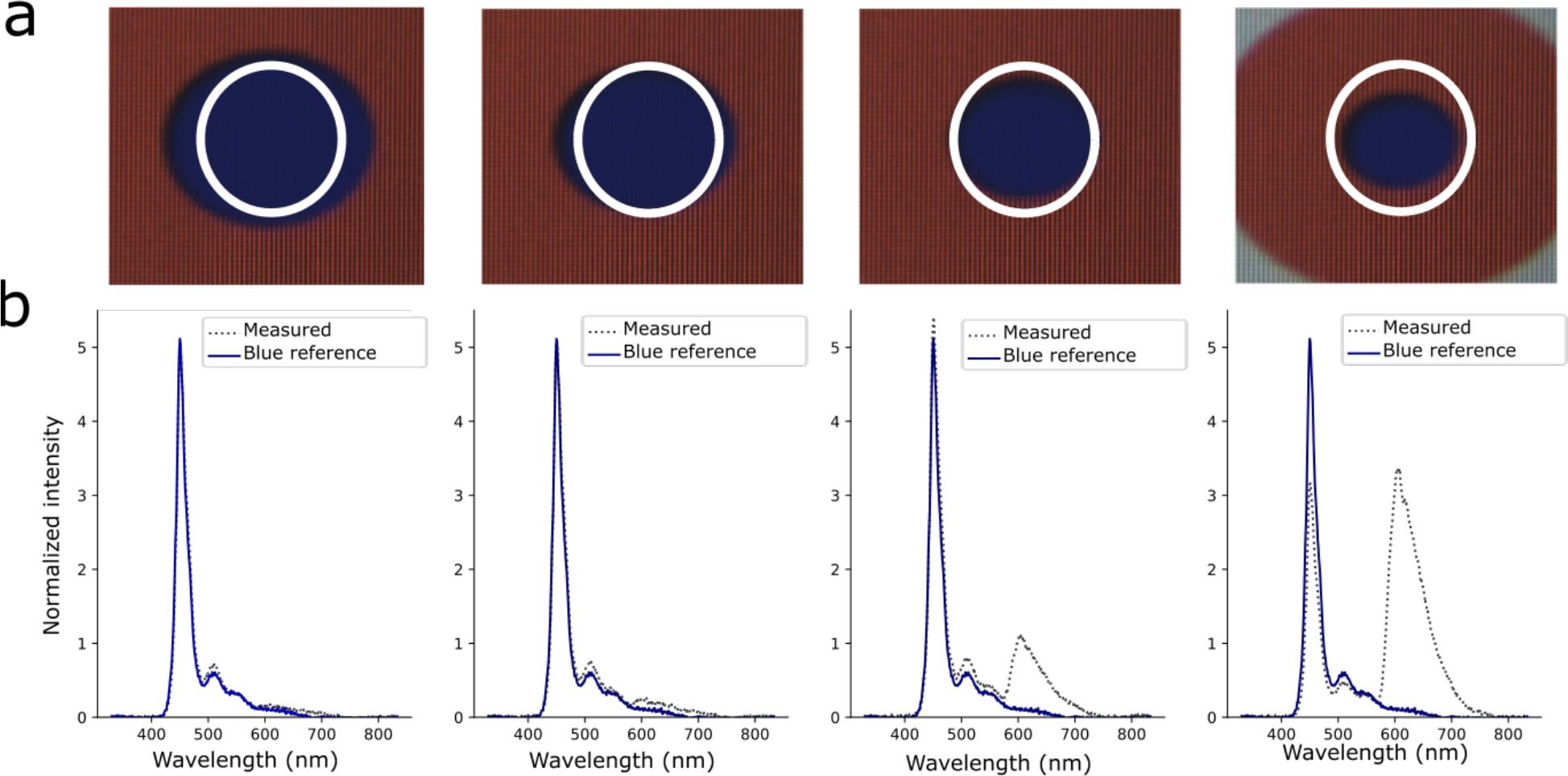
Testing of the overlay of the illumination LED and ROSA for targeted spectroscopy. a) Spectroscopy results compared to blue reference spectrum. b) Visualization of the ROSA overlays. From left to right, the concentricity of the ROSA, depicted with the white circle overlay, was decreased. Results show a decreasing contribution of the blue circle and an increasing contribution from the surrounding red area.

### 3.2 Reflectance acquisition in model eye

Different regions of the model eye were targeted to demonstrate the capacity of the system to acquire reflectance spectra from various targeted regions. Four different regions were targeted – corresponding to blood vessels (A), optic nerve head (B), retina near the optic nerve head (C) and retina far from the optic nerve head (D) – and their spectra were compared (Figure 6).

**Figure 6.**
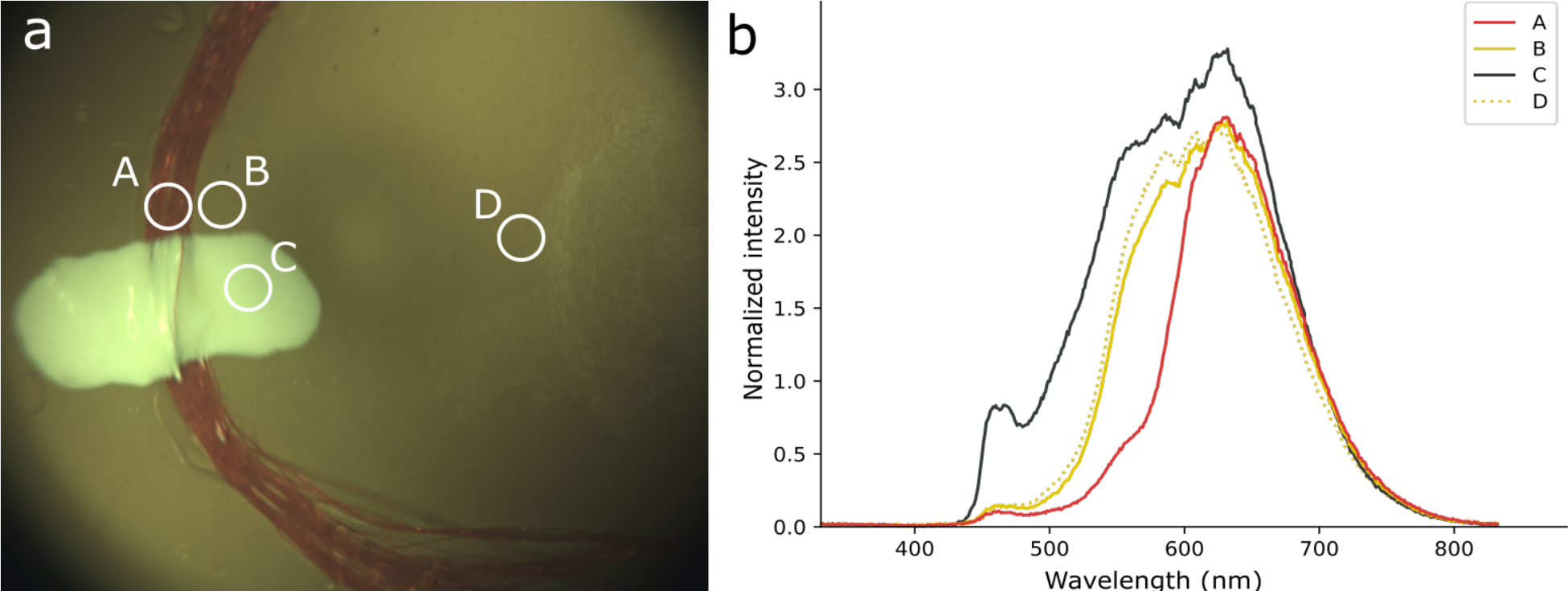
OEMI-7 model eye for targeted reflectance spectroscopy. a) Imaging of the model and the 4 regions of diffuse reflectance spectral acquisitions (A: blood vessels; B: optic nerve head; C: retina near the optic nerve head; D: retina far from the optic nerve head). b) Spectra associated with each region of acquisition.

Regions C and D, which both represent the retina, showed similar reflectance spectra, with only small variations detected. However, clear differences were observed between these regions, region A and region B, corresponding to the blood vessels and the optic nerve head, respectively.

### 3.3 Fluorescence

The model eye used to validate diffuse reflectance acquisition can also be used to perform targeted fluorescence analysis. Thus, different regions were targeted and analyzed for fluorescence on the OEMI-7 (Figure 7). Region A corresponded to a region of high intensity green fluorescence located in the model macula. Region B corresponded to a region located in the model fovea, where no fluorescence signal was visible from the wide field image. Region C corresponded to model blood vessels, with some red fluorescence, located on the simulated retina. Figure 7b shows clear spectral differences between the emitted spectral profiles of regions A and C. Fluorescence signal was only emitted from the targeted region if the targeted region possessed fluorescence capacities, which wasn’t the case for region B.

**Figure 7.**
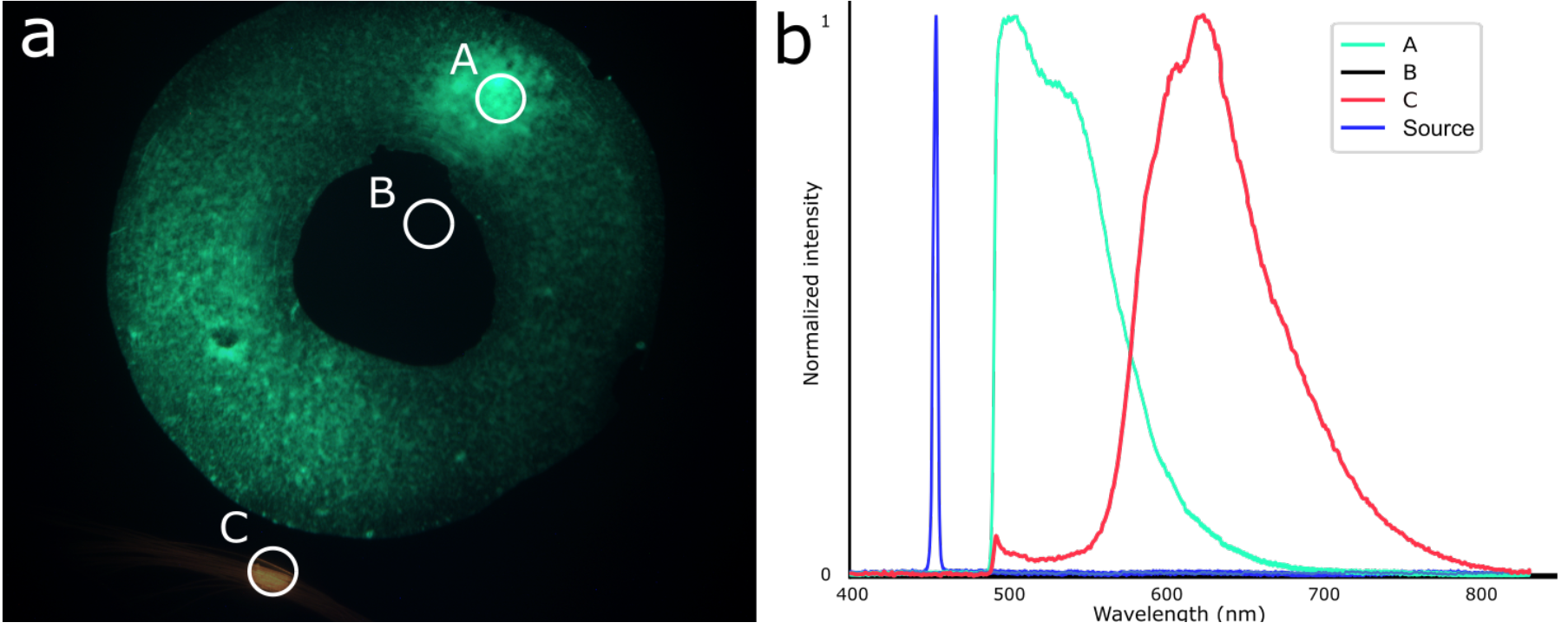
OEMI-7 model eye for fluorescence spectroscopy. a) Fluorescence imaging of the model and identification of the three regions targeted for fluorescence spectral acquisitions. b) Spectra associated with each targeted region (A: green fluorescence; B: no fluorescence; C: red fluorescence) and with the excitation illumination source.

### 3.4 Human eye fundus reflectance

Spectral acquisition was performed in two different regions of the eye fundus – parafovea and optic nerve head, each anatomically different and having different optical properties – of 8 healthy subjects. Figure 8 shows the locations of the acquisitions and the average spectral signatures obtained for the 8 subjects.

**Figure 8.**
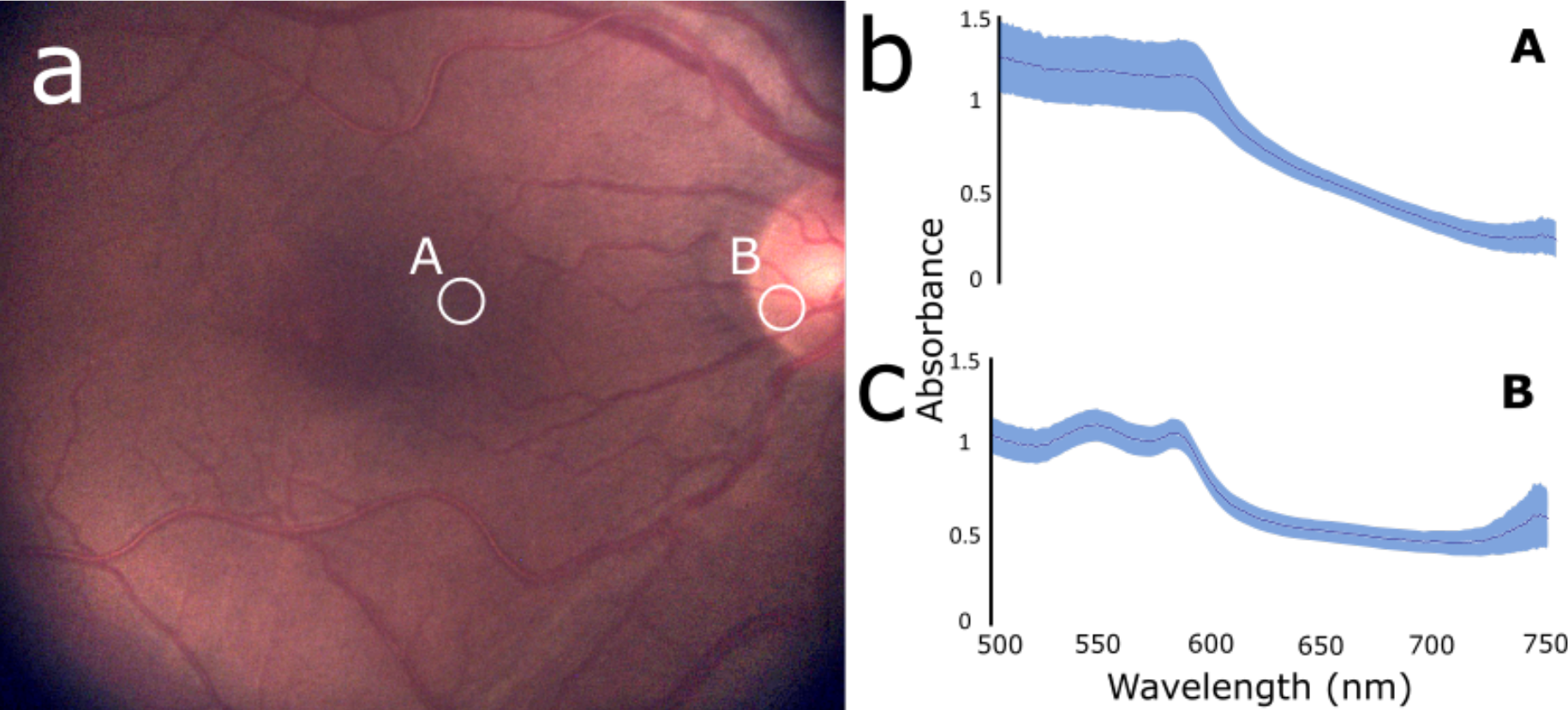
Diffuse reflectance spectral acquisitions on healthy human subjects. a) Locations of the spectral acquisitions performed in healthy subjects – A: parafovea; B: optic nerve head. b) Average absorbance spectra for location A over the 8 subjects. c) Average absorbance spectra for location B over the 8 subjects.

Light intensity was adjusted for every acquisition (subject and location) to cover the whole dynamic range of the spectrometer and improve sensitivity. Five-second acquisitions were made at each location, which corresponds to a total of 13 acquired spectra, with an integration time of 250 ms for the spectrometer and 120 ms for the camera. All acquisitions, taken from the same location, were averaged for every subject. Afterward, the acquisitions for all subjects were average to highlight the differences between regions, represented on Figure 8c.

### 3.5 Oximetry measurements

Assessment of ocular oximetry through the determination of blood oxygen saturation (StO2) at the different locations in the eye fundus of the same 8 healthy subjects was performed as a demonstration of a concrete application stemming from targeted spectral analysis. The StO2 was calculated based on previously described methods [26] for each acquired spectrum and averaged over the acquisition at a specific location for a specific subject. The difference in average StO2 at location A (parafovea) and B (optic nerve head) was compared for each subject, and for the average of the sample population. Results for each subject are reported in Table 1. A Wilcoxon signed-rank test was performed to establish the statistical significance of the difference between the measurements made in the two locations (p = 0.004).

**Table 1.** StO2 values at two locations in the eye fundus of healthy subjects, showing a significant difference between values in the parafovea and the optic disc (Wilcoxon signed-rank test, p-value = 0.004)

| Subject ID | Blood Oxygen Saturation StO<sub>2</sub> (%) |  |  |
| --- | --- | --- | --- |
|  | Parafovea (A) | Optic nerve head (B) | Difference (B-A) |
| 1 | 31.4 | 69.3 | 37.9 |
| 2 | 32.1 | 62.3 | 30.2 |
| 3 | 58.0 | 63.5 | 5.5 |
| 4 | 39.3 | 68.4 | 29.1 |
| 5 | 52.7 | 69.7 | 17.0 |
| 6 | 58.4 | 67.6 | 9.2 |
| 7 | 48.2 | 68.8 | 20.6 |
| 8 | 30.4 | 62.1 | 31.7 |

## 4 Discussion

This work presents a multi-modal system with the ability to concurrently and continuously perform imaging and targeted spectroscopy in the eye fundus. Spectral analysis can be used in the context of reflectance and fluorescence, and has the potential to be applied for a broad range of applications and biomarkers, including ocular oximetry.

In the technology at hand, a pointing LED is used to identify the region for which spectral analysis is to be performed. It was thus important to demonstrate that the projected area of the LED on a surface or on the eye fundus corresponded to the actual ROSA. This was first done in vitro, using reference targets for the evaluation of diffuse reflectance spectra (Figures 4 and 5). Results from alignment and spectral analysis of the first reference target, a high definition screen that uses RGB sources to project custom images, validated that the spectral signal collected by the system came exactly and solely from the region illuminated by the pointing LED (Figure 4). Results from the evaluation of a second reference target, in which blue circles of different sizes were superimposed on red concentric circles to vary the contribution of blue and red colors in the target area, further demonstrated the correspondence of the LED projection with the ROSA (Figure 5). These results also highlighted the sensitivity of the device to the presence of multiple spectral contributors within the ROSA.

It was then important to establish the capacity of the device to distinguish spectral properties of different features of the eye fundus. The OEMI-7 model eye displays features analogous to the optic nerve head, the retina and large blood vessels. Diffuse reflectance spectroscopy performed using this device provided clear, distinguishable spectra for each feature (Figure 6), which corresponded to expected differences in spectral profiles based on color differences observed using an RGB camera. The OEMI-7 model eye was also used to demonstrate how the same technological platform, with minimal modifications in optical layout, was able to acquire defined fluorescence spectra (Figure 7). The spectra for the excitation source, green and red fluorescence regions were clearly distinct from each other. It should also be noted that the fluorescence acquisition from region B, for which there was no fluorescence, demonstrates no spectral signature with negligible noise detected (Figure 7). These in vitro demonstrations of the capacity of the device to provide valuable spectral information from diffuse reflectance and fluorescence at specific targeted locations highlight important features of the technology. The eye fundus is generally regarded as a highly heterogeneous structure, both from a feature (large, blood vessels, microvasculature, macula, fovea, optic nerve head, etc.) and a composition (components, biomarkers, etc.) perspective. For example, biomarkers can be broadly dispersed throughout the tissue (e.g. β-amyloid plaques in the retina of Alzheimer’s patients) [30] or localized in specific regions (e.g. cotton wool spots and hemorrhages for patients with diabetic retinopathy) [31].

The detection of biomarkers from their absorbance or fluorescence spectral signatures is dependent on a few factors, including their distinctive spectral properties and their abundance within the sampling space. There is thus a need for high sensitivity and rapid analysis to avoid some of the limitations encountered in current approaches, such as hyperspectral imaging [32] and VIS-OCT [33]. While extremely useful and informative, these technologies have major drawbacks; the main one being the time required to perform an acquisition, as discussed previously.

Targeted spectroscopy circumvents many of these limitations. It is a technique for fast, continuous spectral acquisition at a specific location of the eye fundus, the ROSA. Wide-range and high-resolution spectra from the ROSA and images of the eye fundus can be acquired within tens of milliseconds. For example, the acquisitions presented in Figure 8 (LED intensity of 1 mW) were performed within 25 ms for the ONH and 175 ms for the parafovea. In this approach, the spectrometer dictates the wavelength range and resolution. In the present configuration, the system can achieve a resolution of 2.3 nm and a spectral response range from 340 to 830 nm; however, these parameters could be broadened, reduced or otherwise modified using a different spectrometer.

In addition, the combination of fast sampling rate with continuous acquisitions opens the door to monitoring retinal changes caused by external stimuli. Morphological variations visible in the images and spectral variations resulting from the response of different absorbers or fluorophores in the retina could easily be quantified by this system. For example, flickering lights has been proven to cause an increase in retinal vascular diameter [34] and retinal and optic nerve blood flow [35]. Quantifying such responses can help identify irregularities in metabolic functions or cellular responses to drugs. Kotliar *et al*. established that neurovascular coupling was altered in Alzheimer’s disease, and that retinal vessel response to flickering light differed in comparison with a control group [36]. Similarly, Song et al. demonstrated that reflectivity from the retinal nerve fiber layer (RNFL) assessed using VIS-OCT was better at distinguishing glaucoma eyes from normal eyes than RNFL thickness measured by OCT [28].

As an extension to the potential applications listed above, targeted spectroscopy could help in the development and assessment of drugs. For example, a number of new drugs targeting optical diseases aim to modulate the oxygenation of ocular tissues [37]. Monitoring the real-time response to such drugs in the specific regions of action could help establish pharmacokinetic parameters and efficacy of treatments. Similarly, tracking the spectral signatures of targeted regions of susceptible tissues during treatment could provide information on the efficacy of the drug.

It seems clear that the capacity to identify a broad range of biomarkers (be they associated with the presence of a compound or a change in structural organization of the tissue) from their spectral signatures opens the door to a wide range of applications. One such example is ocular oximetry, the assessment of blood oxygen saturation in the different tissues of the eye fundus. There is accumulating evidence linking oxygen saturation in specific regions of the eye fundus to a number of diseases, including glaucoma [38]–[43], diabetic retinopathy [38], [39], [44]–[47], age-related macular degeneration [48], occlusions [38], [39], [49], [50], multiple sclerosis [51], and many more. It is thus an increasingly important biomarker.

The evaluation of blood oxygen saturation in the eye has been attempted through methods such as dual-wavelength analysis [52] and hyperspectral imaging [53]. However, in terms of spectral bandwidth and spectral resolution, these techniques do not compete with the sensitivity and accuracy of targeted spectroscopy. This is an important point in the quest for quantitative oxygen measurements, as ultimately spectral resolution and signal-to-noise ratio (SNR) will be the deciding factors in achieving this. Sub-second acquisition speed (usually a byproduct of high SNR) is also very important, as involuntary eye movements called microsaccades typically occur at a frequency of 1.5 Hz, resulting in significantly blurred images and measurements [54]. In their current state, the limited sensitivity of dual-wavelength analysis and hyperspectral imaging only allow for relative assessment of oximetry, and mostly limited to large blood vessels of the eye fundus. Targeted ocular spectroscopy enables assessment in the microvasculature, enabling actual assessment of oximetry and metabolic activity in the tissues.

The results presented in Figure 8 and Table 1 demonstrate how targeted spectroscopy can provide valuable and reproducible measurements of blood oxygen saturation (StO2) in the different regions of the eye fundus. Strong repeatability was observed for multiple acquisitions taken at one location for a given subject. Clear differences were observed in spectra (Figure 8) and StO2 (Table 1) between the parafovea and optic nerve head, with lower oxygen saturation observed in the former. This can be explained mainly by the fact that lower reflected signal in this region results in a decreasing SNR. The fovea is typically the region in the retina with the highest density of photoreceptor cells, leading to higher absorbance and variability compared to the optic nerve head [55]. Moreover, it has been established via OCT-A studies that vascular density decreases as we approach the fovea [56], which explains the decreasing intensity of the hemoglobin spectral components as acquisitions are made further from the ONH. In fact, spectra from the ONH displayed a stronger contribution from hemoglobin components; the nerve fibers, which are the main constituents of the ONH [57] act as diffusers for the main local absorber, which is hemoglobin. The thickness of the different layers of the eye fundus, the contribution of the choroid and the photoreceptor cell density are other factors that can contribute to spectral variations between regions [29].

Another set of potential applications involve fluorescence. Different fluorophores are naturally present in the eye fundus, and they can contribute to diagnosing retinal pathologies. For example, lipofuscin accumulation is associated with age-related macular degeneration and macular dystrophies such as Best and Stargardt disease [58]. Typically, fundus autofluorescence capabilities on a camera enables rough quantification of the amount of molecules present. Our technique enables the user to target specific regions of interest previously identified by wide field fluorescence and then acquire a full spectral profile of the molecules present. This information could be of great use for diagnostic purposes. Feldman *et al*. reported that the emission profile of A2E, a component of RPE, can provide insightful information on the presence of AMD [12].

## 5 Conclusion

The system presented here enables the concurrent and continuous acquisition of images and diffuse reflectance or fluorescence spectra from targeted regions of the eye fundus. This multi- modal approach can provide valuable information on various biomarkers and, by extension, on the physiology and function of specific regions of the eye fundus. This technique has a wide range of potential applications, which include but is not limited to diagnosis and management of retinal diseases, glaucoma, and other conditions that affect the eye. From the acquired spectrum, it is possible to assess the concentrations of different chromophores and fluorophores, such as melanin, hemoglobin and lipofuscin, which are associated with disease progression. By combining fundus imaging with spectral analysis, targeted spectroscopy has the potential to change the way we diagnose and treat eye diseases, and is likely to become an increasingly important tool in eye care in the coming years.

### Disclosures

Nicolas Lapointe, Cléophace Akitegetse, Jasmine Poirier and Maxime Picard are employees of Zilia Inc, which funded this research. Patrick Sauvageau is CEO and co-founder of Zilia Inc. Dominic Sauvageau is CTO and co-founder of Zilia Inc.

## Acknowledgments

This work was supported by funding from the National Research Council of Canada - Industrial Research Assistance Program and the Government of Canada - Scientific Research and Experimental Development Tax Incentive Program.

## Data Availability

Data underlying the results presented in this paper are not publicly available at this time, but may be obtained from the authors upon reasonable request.

## References

1. R. N. Weinreb, T. Aung, and F. A. Medeiros, “The Pathophysiology and Treatment of Glaucoma: A Review,” JAMA, vol. 311, no. 18, p. 1901, May 2014, doi: 10.1001/JAMA.2014.3192.

2. K. Gopalasamy, R. Gayathri, and V. Vishnu Priya, “Age related macular degeneration: A systematic review,” J. Pharm. Sci. Res., vol. 8, no. 6, pp. 416–420, 2016.

3. E. J. Duh, J. K. Sun, and A. W. Stitt, “Diabetic retinopathy: current understanding, mechanisms, and treatment strategies,” JCI Insight, vol. 2, no. 14, Jul. 2017, doi: 10.1172/JCI.INSIGHT.93751.

4. N. K. Archibald, M. P. Clarke, U. P. Mosimann, and D. J. Burn, “The retina in Parkinson’s disease,” Brain, vol. 132, no. 5, pp. 1128–1145, May 2009, doi: 10.1093/BRAIN/AWP068.

5. Querques G, Borrelli E, Sacconi R, De Vitis L, Leocani L, Santangelo R, Magnani G, Comi G, Bandello F. Functional and morphological changes of the retinal vessels in Alzheimer’s disease and mild cognitive impairment. Sci Rep. 2019 Jan 11;9(1):63. doi: 10.1038/s41598-018-37271-6. PMID: 30635610; PMCID: PMC6329813.

6. F. Z. Javaid, J. Brenton, L. Guo, and M. F. Cordeiro, “Visual and Ocular Manifestations of Alzheimer’s Disease and Their Use as Biomarkers for Diagnosis and Progression,” Front. Neurol., vol. 7, Dec. 2016, doi: 10.3389/fneur.2016.00055.

7. Lim JK, Li QX, He Z, Vingrys AJ, Wong VH, Currier N, Mullen J, Bui BV, Nguyen CT. The Eye As a Biomarker for Alzheimer’s Disease. Front Neurosci. 2016 Nov 17;10:536. doi: 10.3389/fnins.2016.00536. PMID: 27909396; PMCID: PMC5112261.

8. Robbins CB, Thompson AC, Bhullar PK, Koo HY, Agrawal R, Soundararajan S, Yoon SP, Polascik BW, Scott BL, Grewal DS, Fekrat S. Characterization of Retinal Microvascular and Choroidal Structural Changes in Parkinson Disease. JAMA Ophthalmol. 2021 Feb 1;139(2):182–188. doi: 10.1001/jamaophthalmol.2020.5730. Erratum in: JAMA Ophthalmol. 2021 Feb 1;139(2):256. PMID: 33355613; PMCID: PMC7758829.

9. M. T. Almonte, P. Capellàn, T. E. Yap, and M. F. Cordeiro, “Retinal correlates of psychiatric disorders.,” Ther. Adv. Chronic Dis., vol. 11, p. 2040622320905215, Mar. 2020, doi: 10.1177/2040622320905215.

10. P. Topcu-Yilmaz, M. Aydin, and B. Cetin Ilhan, “Evaluation of retinal nerve fiber layer, macular, and choroidal thickness in schizophrenia: spectral optic coherence tomography findings,” https://doi.org/10.1080/24750573.2018.1426693, vol. 29, no. 1, pp. 28–33, Jan. 2018, doi: 10.1080/24750573.2018.1426693.

11. A. Ly, L. Nivison-Smith, N. Assaad, and M. Kalloniatis, “Fundus Autofluorescence in Age-related Macular Degeneration,” Optom. Vis. Sci., vol. 94, no. 2, p. 246, Feb. 2017, doi: 10.1097/OPX.0000000000000997.

12. Feldman TB, Yakovleva MA, Larichev AV, Arbukhanova PM, Radchenko AS, Borzenok SA, Kuzmin VA, Ostrovsky MA. Spectral analysis of fundus autofluorescence pattern as a tool to detect early stages of degeneration in the retina and retinal pigment epithelium. Eye (Lond). 2018 Sep;32(9):1440–1448. doi: 10.1038/s41433-018-0109-0. Epub 2018 May 22. PMID: 29786089; PMCID: PMC6137184.

13. X. R. Huang, Y. Zhou, W. Kong, and R. W. Knighton, “Reflectance Decreases before Thickness Changes in the Retinal Nerve Fiber Layer in Glaucomatous Retinas,” Invest. Ophthalmol. Vis. Sci., vol. 52, no. 9, p. 6737, Aug. 2011, doi: 10.1167/IOVS.11-7665.

14. X.-R. Huang, R. W. Knighton, W. J. Feuer, and J. Qiao, “Retinal nerve fiber layer reflectometry must consider directional reflectance,” Biomed. Opt. Express, vol. 7, no. 1, p. 22, Jan. 2016, doi: 10.1364/BOE.7.000022.

15. G. G. Stokes, “On the reduction and oxidation of the colouring matter of the blood,” Proc. R. Soc. Lond., vol. 13, pp. 355–364, Dec. 1864, doi: 10.1098/rspl.1863.0080.

16. D. Van Norren and L. F. Tiemeijer, “Spectral reflectance of the human eye,” Vision Res., vol. 26, no. 2, pp. 313–320, 1986, doi: 10.1016/0042-6989(86)90028-3.

17. F. C. Delori, “Spectrophotometer for noninvasive measurement of intrinsic fluorescence and reflectance of the ocular fundus,” Appl. Opt., vol. 33, no. 31, p. 7439, Nov. 1994, doi: 10.1364/ao.33.007439.

18. M. Hammer, “Imaging spectroscopy of the human ocular fundus in vivo,” J. Biomed. Opt., vol. 2, no. 4, p. 418, 1997, doi: 10.1117/12.285093.

19. V. Diaconu, “Multichannel spectroreflectometry: A noninvasive method for assessment of on-line hemoglobin derivatives,” Appl. Opt., vol. 48, no. 10, pp. D52–D61, Apr. 2009, doi: 10.1364/AO.48.000D52.

20. V. Vucea, P. J. Bernard, P. Sauvageau, and V. Diaconu, “Blood oxygenation measurements by multichannel reflectometry on the venous and arterial structures of the retina,” Appl. Opt., vol. 50, no. 26, pp. 5185–5191, 2011, doi: 10.1364/AO.50.005185.

21. Mordant DJ, Al-Abboud I, Muyo G, Gorman A, Sallam A, Ritchie P, Harvey AR, McNaught AI. Spectral imaging of the retina. Eye (Lond). 2011 Mar;25(3):309–20. doi: 10.1038/eye.2010.222. PMID: 21390065; PMCID: PMC3178323.

22. Desjardins M, Sylvestre JP, Jafari R, Kulasekara S, Rose K, Trussart R, Arbour JD, Hudson C, Lesage F. Preliminary investigation of multispectral retinal tissue oximetry mapping using a hyperspectral retinal camera. Exp Eye Res. 2016 May;146:330–340. doi: 10.1016/j.exer.2016.04.001. Epub 2016 Apr 6. PMID: 27060375.

23. W. R. Johnson, D. W. Wilson, W. Fink, M. Humayun, and G. Bearman, “Snapshot hyperspectral imaging in ophthalmology,” J. Biomed. Opt., vol. 12, no. 1, p. 14036, 2007, doi: 10.1117/1.2434950.

24. J. C. Ramella-Roman and S. A. Mathews, “Spectroscopic measurements of oxygen saturation in the retina,” IEEE J. Sel. Top. Quantum Electron., vol. 13, no. 6, pp. 1697–1703, 2007, doi: 10.1109/JSTQE.2007.911312.

25. Hadoux X, Hui F, Lim JKH, Masters CL, Pébay A, Chevalier S, Ha J, Loi S, Fowler CJ, Rowe C, Villemagne VL, Taylor EN, Fluke C, Soucy JP, Lesage F, Sylvestre JP, Rosa-Neto P, Mathotaarachchi S, Gauthier S, Nasreddine ZS, Arbour JD, Rhéaume MA, Beaulieu S, Dirani M, Nguyen CTO, Bui BV, Williamson R, Crowston JG, van Wijngaarden P. Non-invasive in vivo hyperspectral imaging of the retina for potential biomarker use in Alzheimer’s disease. Nat Commun. 2019 Sep 17;10(1):4227. doi: 10.1038/s41467-019-12242-1. PMID: 31530809; PMCID: PMC6748929.

26. Pi S, Camino A, Cepurna W, Wei X, Zhang M, Huang D, Morrison J, Jia Y. Automated spectroscopic retinal oximetry with visible-light optical coherence tomography. Biomed Opt Express. 2018 Apr 4;9(5):2056–2067. doi: 10.1364/BOE.9.002056. PMID: 29760969; PMCID: PMC5946770.

27. Song W, Zhang S, Kim YM, Sadlak N, Fiorello MG, Desai M, Yi J. Visible Light Optical Coherence Tomography of Peripapillary Retinal Nerve Fiber Layer Reflectivity in Glaucoma. Transl Vis Sci Technol. 2022 Sep 1;11(9):28. doi: 10.1167/tvst.11.9.28. PMID: 36166221; PMCID: PMC9526364.

28. Song W, Shao W, Yi W, Liu R, Desai M, Ness S, Yi J. Visible light optical coherence tomography angiography (vis-OCTA) facilitates local microvascular oximetry in the human retina. Biomed Opt Express. 2020 Jun 30;11(7):4037–4051. doi: 10.1364/BOE.395843. PMID: 33014584; PMCID: PMC7510897.

29. Akitegetse C, Landry P, Robidoux J, Lapointe N, Brouard D, Sauvageau D. Monte-Carlo simulation and tissue-phantom model for validation of ocular oximetry. Biomed Opt Express. 2022 Apr 21;13(5):2929–2946. doi: 10.1364/BOE.458079. PMID: 35774309; PMCID: PMC9203094.

30. D. P. Perl, “Neuropathology of Alzheimer’s Disease,” Mt. Sinai J. Med. N. Y., vol. 77, no. 1, pp. 32–42, 2010, doi: 10.1002/msj.20157.

31. A. A. Alghadyan, “Diabetic retinopathy – An update,” Saudi J. Ophthalmol., vol. 25, no. 2, pp. 99–111, Apr. 2011, doi: 10.1016/j.sjopt.2011.01.009.

32. E. R. Reshef, J. B. Miller, and D. G. Vavvas, “Hyperspectral Imaging of the Retina: A Review,” Int. Ophthalmol. Clin., vol. 60, no. 1, pp. 85–96, 2020, doi: 10.1097/IIO.0000000000000293.

33. X. Shu, L. J. Beckmann, and H. F. Zhang, “Visible-light optical coherence tomography: a review,” J. Biomed. Opt., vol. 22, no. 12, p. 121707, Dec. 2017, doi: 10.1117/1.JBO.22.12.121707.

34. A. E. Felder, J. Wanek, N. P. Blair, and M. Shahidi, “Inner Retinal Oxygen Extraction Fraction in Response to Light Flicker Stimulation in Humans,” Invest. Ophthalmol. Vis. Sci., vol. 56, no. 11, pp. 6633–6637, Oct. 2015, doi: 10.1167/iovs.15-17321.

35. Palkovits S, Lasta M, Told R, Schmidl D, Werkmeister R, Cherecheanu AP, Garhöfer G, Schmetterer L. Relation of retinal blood flow and retinal oxygen extraction during stimulation with diffuse luminance flicker. Sci Rep. 2015 Dec 17;5:18291. doi: 10.1038/srep18291. PMID: 26672758; PMCID: PMC4682144.

36. Kotliar K, Hauser C, Ortner M, Muggenthaler C, Diehl-Schmid J, Angermann S, Hapfelmeier A, Schmaderer C, Grimmer T. Altered neurovascular coupling as measured by optical imaging: a biomarker for Alzheimer’s disease. Sci Rep. 2017 Oct 10;7(1):12906. doi: 10.1038/s41598-017-13349-5. PMID: 29018233; PMCID: PMC5635105.

37. J. Gibson, “Clinical aspects of retinal oximetry,” Acta Ophthalmol. (Copenh.), vol. 90, no. s249, p. 0, Dec. 2018, doi: 10.1111/j.1755-3768.2012.2816.x.

38. J. Boeckaert, E. Vandewalle, and I. Stalmans, “Oximetry: recent insights into retinal vasopathies and glaucoma.,” Bull. Soc. Belge Ophtalmol., vol. 319, no. 319, pp. 75–83, 2012.

39. S. H. Hardarson, “Retinal Oximetry,” Acta Ophthalmol. (Copenh.), vol. 91, no. thesis2, pp. 1–47, Dec. 2013, doi: 10.1111/aos.12086.

40. Olafsdottir OB, Vandewalle E, Abegão Pinto L, Geirsdottir A, De Clerck E, Stalmans P, Gottfredsdottir MS, Kristjansdottir JV, Van Calster J, Zeyen T, Stefánsson E, Stalmans I. Retinal oxygen metabolism in healthy subjects and glaucoma patients. Br J Ophthalmol. 2014 Mar;98(3):329–33. doi: 10.1136/bjophthalmol-2013-303162. Epub 2014 Jan 8. PMID: 24403567.

41. Tobe LA, Harris A, Schroeder A, Gerber A, Holland S, Amireskandari A, Kim NJ, Siesky B. Retinal oxygen saturation and metabolism: how does it pertain to glaucoma? An update on the application of retinal oximetry in glaucoma. Eur J Ophthalmol. 2013 Jul-Aug;23(4):465- 72. doi: 10.5301/ejo.5000289. Epub 2013 Apr 18. PMID: 23640511.

42. Vandewalle E, Abegão Pinto L, Olafsdottir OB, De Clerck E, Stalmans P, Van Calster J, Zeyen T, Stefánsson E, Stalmans I. Oximetry in glaucoma: correlation of metabolic change with structural and functional damage. Acta Ophthalmol. 2014 Mar;92(2):105–10. doi: 10.1111/aos.12011. Epub 2013 Jan 17. PMID: 23323611.

43. D. J. Mordant, I. Al-Abboud, G. Muyo, A. Gorman, A. R. Harvey, and A. I. McNaught, “Oxygen saturation measurements of the retinal vasculature in treated asymmetrical primary open-angle glaucoma using hyperspectral imaging,” Eye, vol. 28, no. 10, pp. 1190–1200, 2014, doi: 10.1038/eye.2014.169.

44. C. M. Jørgensen, S. H. Hardarson, and T. Bek, “The oxygen saturation in retinal vessels from diabetic patients depends on the severity and type of vision-threatening retinopathy,” Acta Ophthalmol. (Copenh.), vol. 92, no. 1, pp. 34–39, Feb. 2014, doi: 10.1111/aos.12283.

45. S. H. Hardarson and E. Stefánsson, “Retinal oxygen saturation is altered in diabetic retinopathy,” Br. J. Ophthalmol., vol. 96, no. 4, pp. 560–563, Apr. 2012, doi: 10.1136/bjophthalmol-2011-300640.

46. A. Guduru, T. G. Martz, A. Waters, A. V. Kshirsagar, and S. Garg, “Oxygen saturation of retinal vessels in all stages of diabetic retinopathy and correlation to ultra-wide field fluorescein angiography,” Invest. Ophthalmol. Vis. Sci., vol. 57, no. 13, pp. 5278–5284, Oct. 2016, doi: 10.1167/iovs.16-20190.

47. Hammer M, Vilser W, Riemer T, Mandecka A, Schweitzer D, Kühn U, Dawczynski J, Liemt F, Strobel J. Diabetic patients with retinopathy show increased retinal venous oxygen saturation. Graefes Arch Clin Exp Ophthalmol. 2009 Aug;247(8):1025–30. doi: 10.1007/s00417-009-1078-6. Epub 2009 Apr 29. PMID: 19404666.

48. A. Geirsdottir, S. H. Hardarson, O. B. Olafsdottir, and E. Stefánsson, “Retinal oxygen metabolism in exudative age-related macular degeneration,” Acta Ophthalmol. (Copenh.), vol. 92, no. 1, pp. 27–33, Dec. 2014, doi: 10.1111/aos.12294.

49. T. H. Williamson, J. Grewal, B. Gupta, B. Mokete, M. Lim, and C. H. Fry, “Measurement of PO2 during vitrectomy for central retinal vein occlusion, a pilot study,” Graefes Arch. Clin. Exp. Ophthalmol., vol. 247, no. 8, pp. 1019–1023, Dec. 2009, doi: 10.1007/s00417-009-1072-z.

50. Yoneya S, Saito T, Nishiyama Y, Deguchi T, Takasu M, Gil T, Horn E. Retinal oxygen saturation levels in patients with central retinal vein occlusion. Ophthalmology. 2002 Aug;109(8):1521–6. doi: 10.1016/s0161-6420(02)01109-0. PMID: 12153805.

51. Stefánsson E, Olafsdottir OB, Einarsdottir AB, Eliasdottir TS, Eysteinsson T, Vehmeijer W, Vandewalle E, Bek T, Hardarson SH. Retinal Oximetry Discovers Novel Biomarkers in Retinal and Brain Diseases. Invest Ophthalmol Vis Sci. 2017 May 1;58(6):BIO227-BIO233. doi: 10.1167/iovs.17-21776. PMID: 28810002.

52. A. Geirsdottir, O. Palsson, S. H. Hardarson, O. B. Olafsdottir, J. V. Kristjansdottir, and E. Stefánsson, “Retinal vessel oxygen saturation in healthy individuals,” Invest. Ophthalmol. Vis. Sci., vol. 53, no. 9, pp. 5433–5442, 2012, doi: 10.1167/iovs.12-9912.

53. Kaluzny J, Li H, Liu W, Nesper P, Park J, Zhang HF, Fawzi AA. Bayer Filter Snapshot Hyperspectral Fundus Camera for Human Retinal Imaging. Curr Eye Res. 2017 Apr;42(4):629–635. doi: 10.1080/02713683.2016.1221976. Epub 2016 Oct 21. PMID: 27767345; PMCID: PMC5389919.

54. R. G. Alexander, S. L. Macknik, and S. Martinez-Conde, “Microsaccade Characteristics in Neurological and Ophthalmic Disease.,” Front. Neurol., vol. 9, p. 144, 2018, doi: 10.3389/fneur.2018.00144.

55. R. Legras, A. Gaudric, and K. Woog, “Distribution of cone density, spacing and arrangement in adult healthy retinas with adaptive optics flood illumination,” PLoS ONE, vol. 13, no. 1, 2018, doi: 10.1371/journal.pone.0191141.

56. You QS, Chan JCH, Ng ALK, Choy BKN, Shih KC, Cheung JJC, Wong JKW, Shum JWH, Ni MY, Lai JSM, Leung GM, Cheung CMG, Wong TY, Wong IYH. Macular Vessel Density Measured With Optical Coherence Tomography Angiography and Its Associations in a Large Population-Based Study. Invest Ophthalmol Vis Sci. 2019 Nov 1;60(14):4830–4837. doi: 10.1167/iovs.19-28137. PMID: 31747685.

57. J. Salazar J, I. Ramírez A, De Hoz R, Salobrar-Garcia E, Rojas P, A. Fernández-Albarral J, et al. Anatomy of the Human Optic Nerve: Structure and Function [Internet]. Optic Nerve. IntechOpen; 2019. Available from: http://dx.doi.org/10.5772/intechopen.79827

58. G. L. Wing, G. C. Blanchard, and J. J. Weiter, “The topography and age relationship of lipofuscin concentration in the retinal pigment epithelium.,” Invest. Ophthalmol. Vis. Sci., vol. 17, no. 7, pp. 601–607, Jul. 1978.

