## Supplemental Figure S1 for "Targeted spectroscopy in the eye fundus"

### Supplemental Material

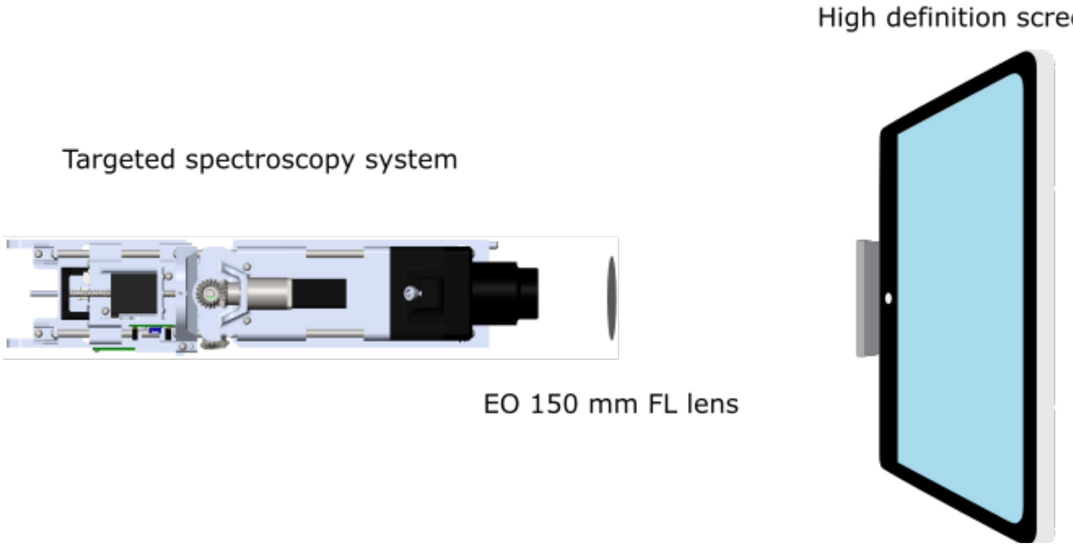


**Figure S1**. Configuration for the validation of the region of spectral acquisition. The targeted spectroscopy system is placed in front of a high-definition screen. A lens is placed in between to relay the image onto the sensor. The screen is used to display different and customizable targets to validate the system.
